# The value of corticospinal excitability and intracortical inhibition in predicting motor skill improvement driven by action observation

**DOI:** 10.1101/2021.10.07.463481

**Authors:** Arturo Nuara, Chiara Bazzini, Pasquale Cardellicchio, Emilia Scalona, Doriana De Marco, Giacomo Rizzolatti, Maddalena Fabbri-Destro, Pietro Avanzini

## Abstract

**BACKGROUND AND OBJECTIVE:** Action observation can sustain motor skill improvement. At the neurophysiological level, action observation affects the excitability of the motor cortices, as measured by transcranial magnetic stimulation. However, whether the cortical modulations induced by action observation may explain the amount of motor improvement driven by action observation training (AOT) remains to be addressed.

**METHODS:** We conducted a two-phase study involving 40 volunteers. First, we assessed the effect of action observation on corticospinal excitability (amplitude of motor evoked potentials), short-interval intracortical inhibition, and transcallosal inhibition (ipsilateral silent period). Subsequently, a randomized-controlled design was applied, with AOT participants asked to observe and then execute, as quickly as possible, a right-hand dexterity task six consecutive times, whereas controls had to observe a no-action video before performing the same task.

**RESULTS:** AOT participants showed greater performance improvement relative to controls. The amount of improvement in the AOT group was predicted by the amplitude of corticospinal modulation during action observation and even more by the amount of intracortical inhibition induced by action observation. Importantly, these relations were found specifically for the AOT group and not for the controls.

**CONCLUSIONS:** In this study, we identified the neurophysiological signatures associated with, and potentially sustaining, the outcome of AOT. Intracortical inhibition driven by action observation plays a major role. These findings elucidate the cortical mechanisms underlying AOT efficacy and open to predictive assessments for the identification of potential responders to AOT, informing the rehabilitative treatment individualization.

## Introduction

Action observation plays a key role in promoting neuroplasticity processes underlying motor learning [1–4]. Following this principle, a motor training approach grounded in the alternation of action observation and execution (i.e., action observation training [AOT]) has been developed to promote the acquisition and recovery of motor abilities [3].

The transformation of sensory representations of others’ actions into one’s motor representation (i.e., mirror mechanism [5]) represents a key process by which AOT may lead to behavioral effects. Indeed, at the neurophysiological level, action observation affects the excitability of the motor system, which can be measured by transcranial magnetic stimulation (TMS), corticospinal excitability assessment [6–8], intracortical inhibition [7, 9, 10], and transcallosal inhibition [11] (see also Naish et al. [12]). However, whether these corticomotor modulations evoked by action observation explain the individual amount of motor improvement driven by AOT remains to be addressed.

For this purpose, we evaluated, via TMS, the effects of action observation on (1) corticospinal excitability (motor evoked potentials [MEPs]), (2) short-interval intracortical inhibition (sICI), and (3) transcallosal inhibition (ipsilateral silent period [iSP]) in 40 healthy participants. Subsequently, we administered either an AOT or, as a control, motor training with observation of non-action videos. Finally, we assessed the capacity of each neurophysiological marker to predict AOT outcomes. The identification of predictors sheds light on the cortical mechanisms underlying AOT efficacy and sets the premises for developing assessments aimed at identifying the best candidates for AOT.

## Methods

### Participants

Forty subjects (10 males and 30 females, mean age 36 ± SD 9 years [range 22–61 years]) were recruited for the study. All subjects were right-handed, as assessed by the Edinburgh Handedness Inventory [13]. None of them had any history of neurological/psychiatric diseases or contraindications to TMS administration [14]. Participants were informed about the experimental procedures and gave their written consent according to the Helsinki Declaration. The experiment was approved by the local ethical committee “Area Vasta Emilia Nord” (n. 10084, 13.03.2018).

In the next paragraphs, we will detail the experimental design, which is graphically summarized in Figure 1.

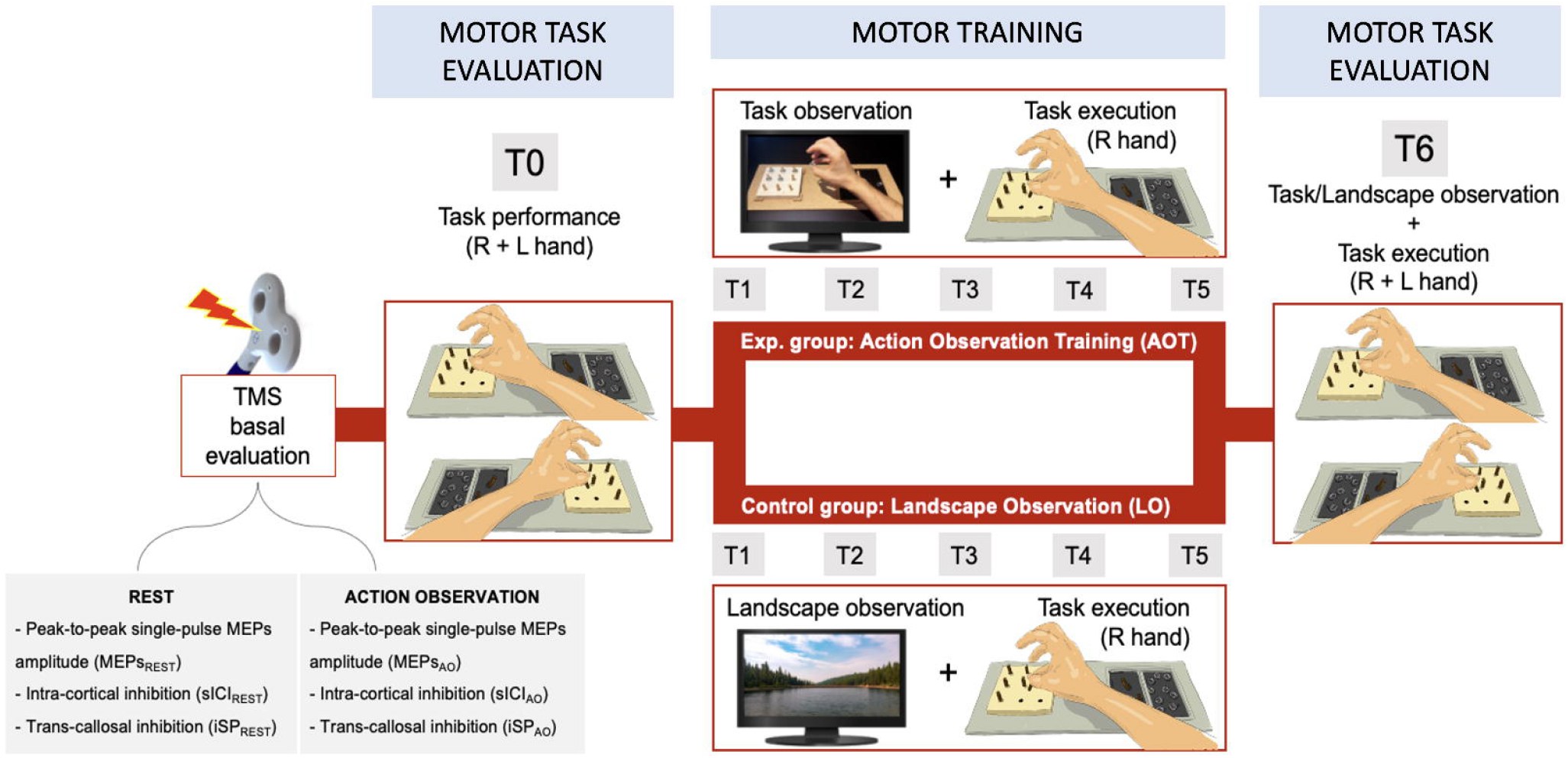

### Baseline evaluation

TMS was delivered by a figure-of-eight coil (70 mm) connected to a Magstim BiStim stimulator (Magstim, Whitland, UK) and combined with electromyographic (EMG) measurements to assess MEPs. TMS was applied to the scalp, with the coil handle rotated 45° from the sagittal plane. Before the experimental session, the optimal stimulation location (hotspot) corresponding to the right first digital interosseous (R-FDI) was determined. The hotspot was defined as the scalp location providing the highest peak-to-peak MEP amplitude in the relaxed R-FDI averaged over five consecutive stimuli. The coil position and orientation were coregistered to a brain template obtained from individual head landmarks (nasion, ears, scalp surface) using an optoelectronic neuronavigation system (visor 2, ANT Neuro, Netherlands).

EMG signals from the R-FDI muscle and the left opponens pollicis (L-OP) were continuously recorded using surface Ag–AgCl electrodes. The EMG signal was amplified (×1000) using a CED1902 amplifier (Cambridge Electronic Design), sampled at 2.5 kHz, filtered with an analogical online band-pass (20–250 Hz) and a notch (50 Hz) filter, and acquired with CED Micro 1401 interfaced with Spike2 software (Cambridge Electronic Design). An additional channel containing digital markers of the TMS trigger was integrated into the same EMG file. The data were stored for subsequent analyses.

The corticomotor excitability was assessed with the following TMS parameters:

a. The resting motor threshold (RMT), defined as the lowest stimulator output intensity capable of inducing MEPs greater than 50 μV peak-to-peak amplitude in relaxed R-FDI in at least 5 of 10 trials [15].
b. Peak-to-peak amplitude of MEPs elicited in the resting R-FDI by single-pulse TMS (120% RMT intensity).
c. sICI was obtained from a paired-pulse TMS protocol [16]. A subthreshold conditioning stimulus was delivered at 80% of the RMT and at an interstimulus interval of 3 ms before a suprathreshold, conditioned, test stimulus (120% RMT). Both stimuli were delivered by the same coil in the same scalp position. sICI was expressed as the percentage decrease of MEP amplitude relative to the single-pulse TMS condition, according to the following formula:

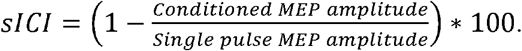
d. iSP was acquired by delivering single-pulse TMS to the right opponens pollicis hotspot (obtained with a procedure like that described for the R-FDI muscle) while the participant maintained a maximal contraction of the L-OP.

The iSP parameters were computed from the rectified traces of the L-OP EMG. The iSP onset was defined as the point at which EMG activity decreased (minimum duration 10 ms) of at least 2 standard deviations relative to the baseline (60-10 ms prestimulus). The iSP offset was defined as the first point after iSP onset at which the EMG activity regained the baseline value. The iSP_AREA_ was defined as the EMG area between iSP offset and iSP onset, while Baseline_AREA_ as the EMG area between 60-10 ms before the TMS stimulus [17]. We then calculated the iSP amount according to the formula:

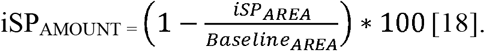

Subjects performed the experiment seated in a comfortable armchair and in front of a 17-inch LCD computer monitor (1024 × 768 pixels) placed 60 cm from their frontal plane. First, the abovementioned TMS parameters were measured during the continuous observation of a black screen with a white cross in its center (REST). Three separate sessions lasting two minutes were administered, one for each specific TMS parameter (standard MEPs, sICI, and iSP). While subjects were asked to keep their upper limbs relaxed during standard MEPs and sICI assessments, during the iSP assessment, they were requested to start the voluntary contraction upon the verbal request of the experimenter, who controlled and jittered the delivery of TMS pulses. Within each session, 15 TMS pulses were administered.

After the REST protocol, the same parameters listed above were estimated during action observation. In this protocol, subjects were asked to carefully observe 24 video clips depicting reach-to-grasp actions toward different objects. Each video, showing a pinch- or tri-digital grasp, represented the action from a first-person perspective and lasted 3.5 s. An intertrial (2 s, black background) was interposed between the videos. The overall action observation trial duration was about 2 minutes, in line with the resting condition. During the iSP assessment, subjects were requested to start the voluntary contraction at each action onset and to relax during the intertrial. Within each session, in 15 of the 24 videos, TMS was randomly delivered 200 ms prior to hand-object contact. Such a latency has been previously shown as the timepoint providing the maximal MEP amplitude [8, 19]. Considering potential repetition suppression phenomena related to the TMS series [20], the protocol sequence was randomized across participants.

### Motor training

A modified version of the Nine Hole Peg Test (mNHPT)—a quantitative test of upper extremity function [21]—was adopted to assess motor performance. Previous studies have shown that the performance of the standard NHPT strongly depends on frontoparietal network functioning [22] [23]. Moreover, NHPT performance improves with repetition over time [24], denoting the test’s suitability as a motor learning endpoint. At baseline (T0), both the right and left hands were tested (see Figure 1, task performance). Participants were seated at a table hosting a woodblock with nine empty holes on one side and a small container on the other. The latter was further split into two parts holding nine pegs and nine nuts, respectively.

On a start cue, subjects had to pick up the nine pegs one at a time as quickly as possible and put them in the nine holes according to a preestablished order (left to right, top to bottom). After placing the pegs in the holes, they had to apply the nuts in correspondence with each peg, following the same insertion order. Finally, they had to remove the nuts and pegs as quickly as possible—one at a time, placing them back into the proper container. Noteworthy, subjects were asked to adopt a first–fifth pinch grasp (thumb–little finger) throughout the task. This constraint, as well as the adding of the nuts, was introduced in the modified version of the test to increase task difficulty, thus delaying the performance “ceiling effect.” The task was video-recorded and scored offline. The time required to perform the mNHPT was selected as the primary endpoint. In addition, errors, defined as placing, sequence, or hand-posture inaccuracies, were registered.

Subjects were randomized into two groups (see upper and lower strips in Figure 1). Participants belonging to the experimental (AOT) group (n = 20) were asked to observe a video clip showing a correct right-hand execution of the mNHPT (duration 1:16 min) and then to execute it as quickly and accurately as possible. This observation-execution combination was repeated six consecutive times (namely, T1–T6). The last trial (T6) also included left-hand mNHPT execution, thus allowing a direct before and after training comparison of both hands’ performance. Participants in the control (landscape-observation) group (n = 20) followed the same procedure, except the content of the video clip preceding the mNHPT execution depicted a landscape.

The time required to perform the mNHPT was recorded at each timepoint. The percentage decrease of total time relative to T0 (in other words, the increased speed) was computed. The T0– T6 percentage difference in right-hand mNHPT execution speed was set as the main behavioral endpoint. Secondary endpoints included T0–T6 left-hand improvement in mNHPT execution speed.

### Data analysis

The effect of action observation on motor cortex excitability was assessed by comparing the TMS parameters (MEPs, sICI, iSP) between the *rest* and *action observation* protocols by means of direct, nonparametric contrasts (Wilcoxon test). The choice of nonparametric tests was due to the absence of normality assumption.

Beyond investigating the modulations induced by action observation at the population level, we also moved to the individual level, thus computing the ratio between action observation and REST protocols for each of the TMS parameters:

a. MEPs_(AO)_/MEPs_(REST)_
b. sICI_(AO)_/sICI_(REST)_
c. iSP_(AO)_/iSP_(REST)_

Mixed ANOVA was applied to the right-hand mNHPT speed increase, considering TIME as a within-subject factor and GROUP as a between-subjects factor. As T0 served as a baseline for individual data, six levels were included in TIME (T1–T6). Planned comparisons were made using independent sample, two-tailed t-tests, limited to the comparison between groups at each timepoint. A preliminary analysis was conducted to evaluate the correlation between basal TMS parameters estimated at rest (MEP amplitude, sICI, and iSP) and motor improvement in the whole population, regardless of the specific group, using Spearman’s rank correlation.

Subsequently, the correlation between the basal neurophysiological features assessing left-hemisphere excitability modulation by action observation (MEPs_(AO)_/MEPs_(REST)_, sICI_AO_/sICI_REST_) and motor outcomes (right- and left-hand T0–T6 mNHPT improvement) was separately evaluated in each group using Spearman’s rank correlation. iSP_AO_/iSP_REST_ was correlated with left-hand T0– T6 mNHPT improvement only.

In case of significant correlations, the capability of the neurophysiological modulations induced by action observation to predict motor improvement in subjects undergoing AOT was tested by applying a linear regression model. In case of multiple significances, multiple linear regression models were also applied to evaluate the cross-talk between individual regressors. For each significant regression, a Bayesian factor (*BF*_*1*|*0*_) was computed to quantify the evidence in favor of the alternative hypothesis (i.e., the neurophysiological feature *predicts* motor outcome) relative to the null hypothesis (i.e., the neurophysiological feature *does not predict* motor outcome). Despite being widely adopted and easy to interpret as a motor learning endpoint, the mere difference between T0 and T6 does not account for the temporal dynamics of the learning process. Indeed, regardless of the T6 performance, the learning curve at T6 could exhibit higher/lower slopes. Thus, for each subject, we applied a regression model to fit the timewise perxformances into a logarithmic curve defined by the following equation: *y* = *A* * log(*bx*), where *x* indicates the trial number and the *A* coefficient indexes the slope of the curve. In the case of significant regression, the *A* coefficient can be regarded as a time-independent index of *motor learning drive*. Significant results would extend the validity of timepoint-specific observations to a global, time-independent dynamic. Then *A* values were compared between groups by direct contrast (independent samples, two-tailed t-test), and following the same statistical procedures described above, a linear regression was performed against baseline neurophysiological variables.

## Results

### Participants’ compliance and safety

All the experimental procedures were well tolerated. In particular, no side effects related to TMS administration were reported. One subject did not complete the experimental procedures, even though she completed the baseline neurophysiological evaluation. Concerning sICI, one control subject was excluded from the neurophysiological evaluation due to the triggering system malfunctioning.

### Participants’ baseline features

An independent samples t-test did not detect significant between-group differences in baseline behavioral and neurophysiological features (right mNHPT: t[39] = −0.762, p = 0.45; left mNHPT: t[39] = −0.240, p = 0.81; RMT: t[39] = 0.115, p = 0.91; MEP amplitude: t[39] = 0.66, p = 0.512; sICI: t[39] = 1.870, p = 0.70; iSP: t[39] = 0.708, p = 0.48).

### Effect of action observation on corticomotor excitability

Single-pulse MEPs elicited during action observation were significantly higher than during the resting condition (1.46 ± 1.06 vs 1.65 ± 1.09 mV, Z[40] = 2.460, p = 0.014, see Figure 2, Panel A), indicating an average facilitation effect of 13%. Although the overall effect was significant, a consistent variability emerged at the single-subject level; 14 out of 40 subjects (35%) showed a decrease in MEP amplitude during action observation.

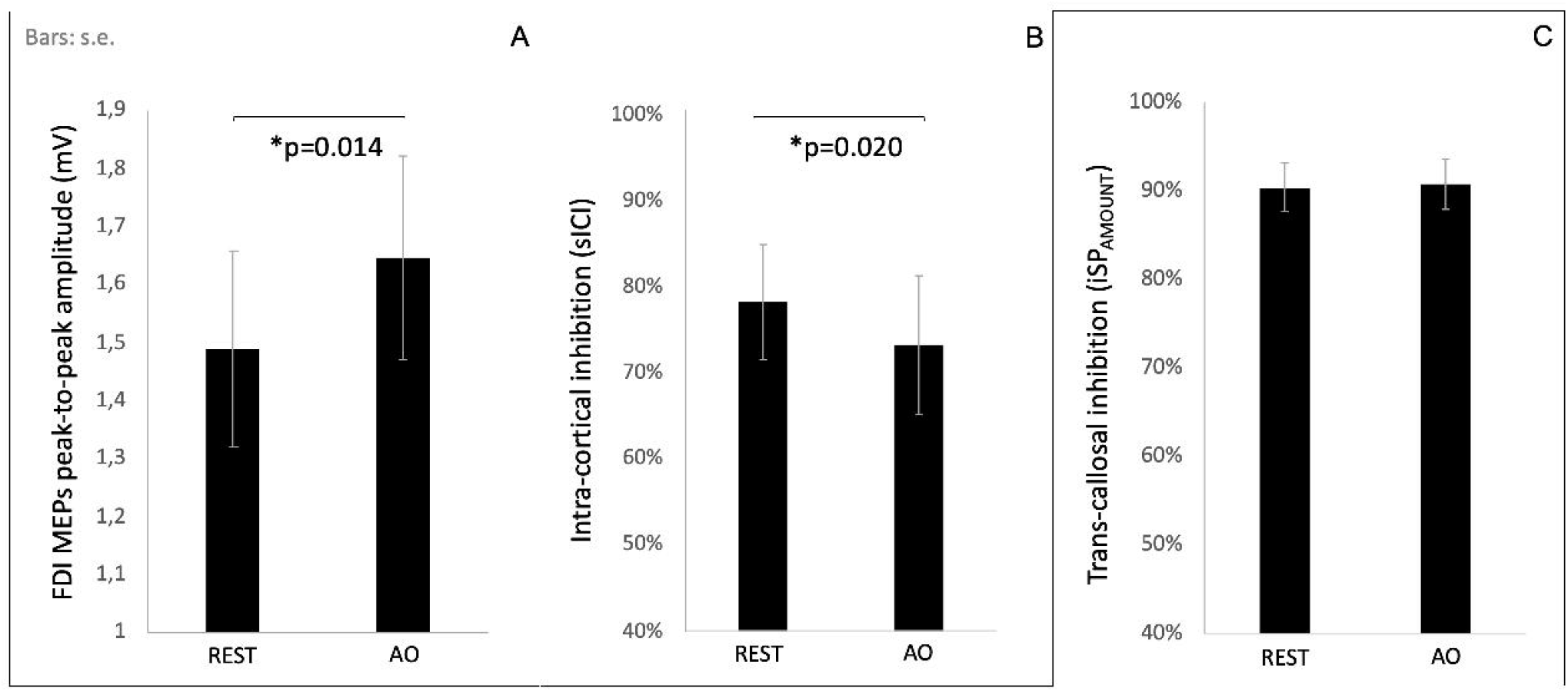

Action observation induced a significant decrease in sICI (78.25% ± 20.95% vs. 73.26% ± 25.00%, Z[39] = −1.619, p = 0.02, Figure 2, Panel B). Even in this case, despite the overall significant decrease, 13 out of the 39 participants (32%) displayed an increase in sICI during action observation. A little overlap (n = 1) was observed between the 14 MEP suppressors and the 13 sICI enhancers. No significant change was observed comparing ISP amount at rest vs. during action observation (90.47% ± 27.35 vs 90.88% ± 28.63 p = 0.35, see Figure 2, Panel C).

### Effect of action observation on motor improvement

Repeated measure ANOVA showed that both TIME (F [5, 185], p<0.001) and GROUP (F [1, 37], p<0.001) factors had a significant effect on mNHPT speed. Planned contrasts indicated that subjects undergoing AOT had greater improvement than controls since the first execution and throughout all the timepoints (see Figure 3, Panel A). No significant time*group interaction was found. The main motor outcome—that is, overall improvement at T6 in right-hand mNHPT speed—was greater in AOT subjects in comparison to controls (27.67% ± 6.4 vs. 19.01% ± 3.1; t[39] = −5.362; p<0.001; η^2^ = 0.437). There were similar findings regarding left-hand mNHPT performance at T6 (controls 14.55% ± 7.95 vs. AOT 20.01% ± 7.16; t[39] = −2.288; p = 0.028; η^2^ = 0.124; see Figure 4, Panel A).

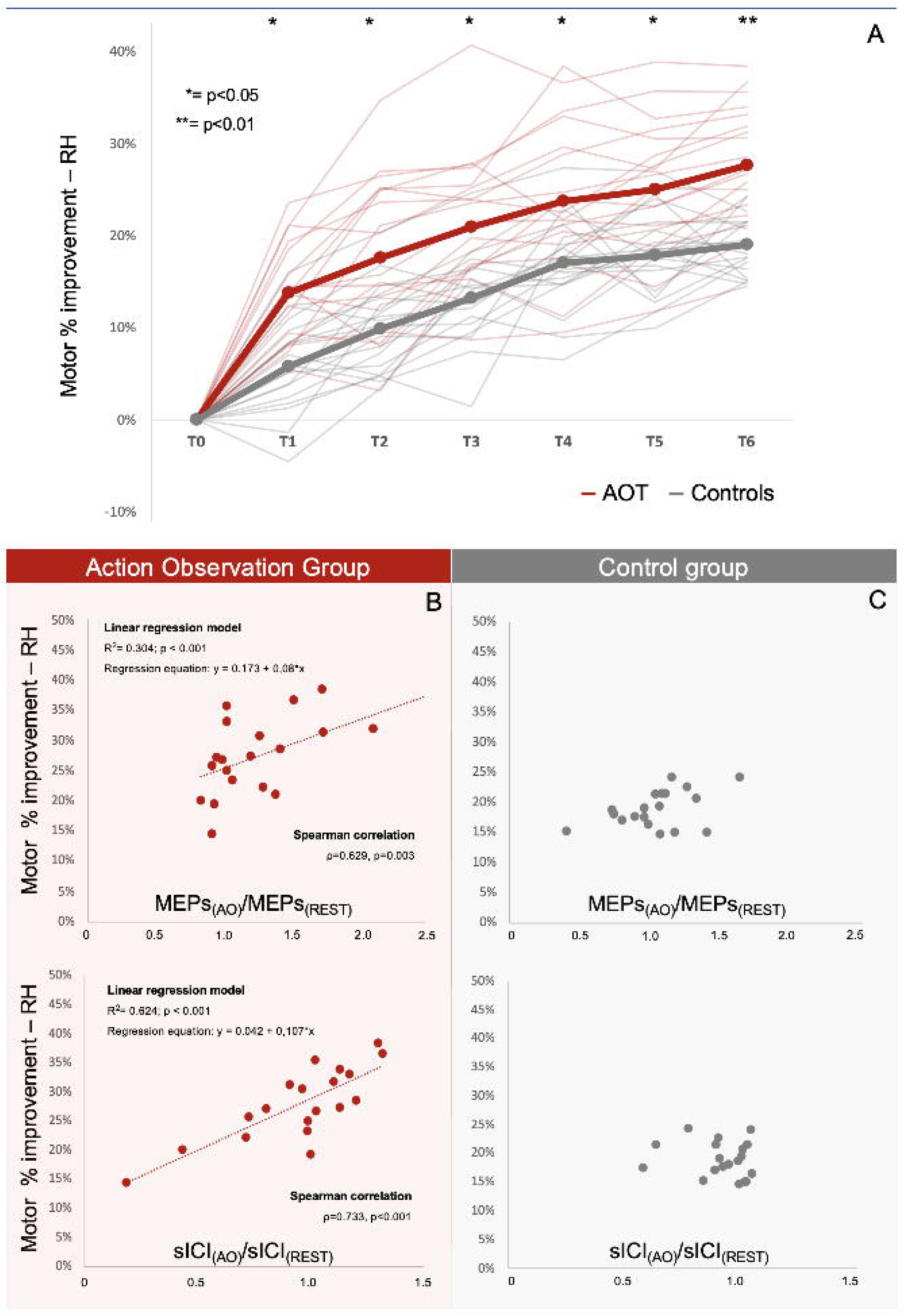
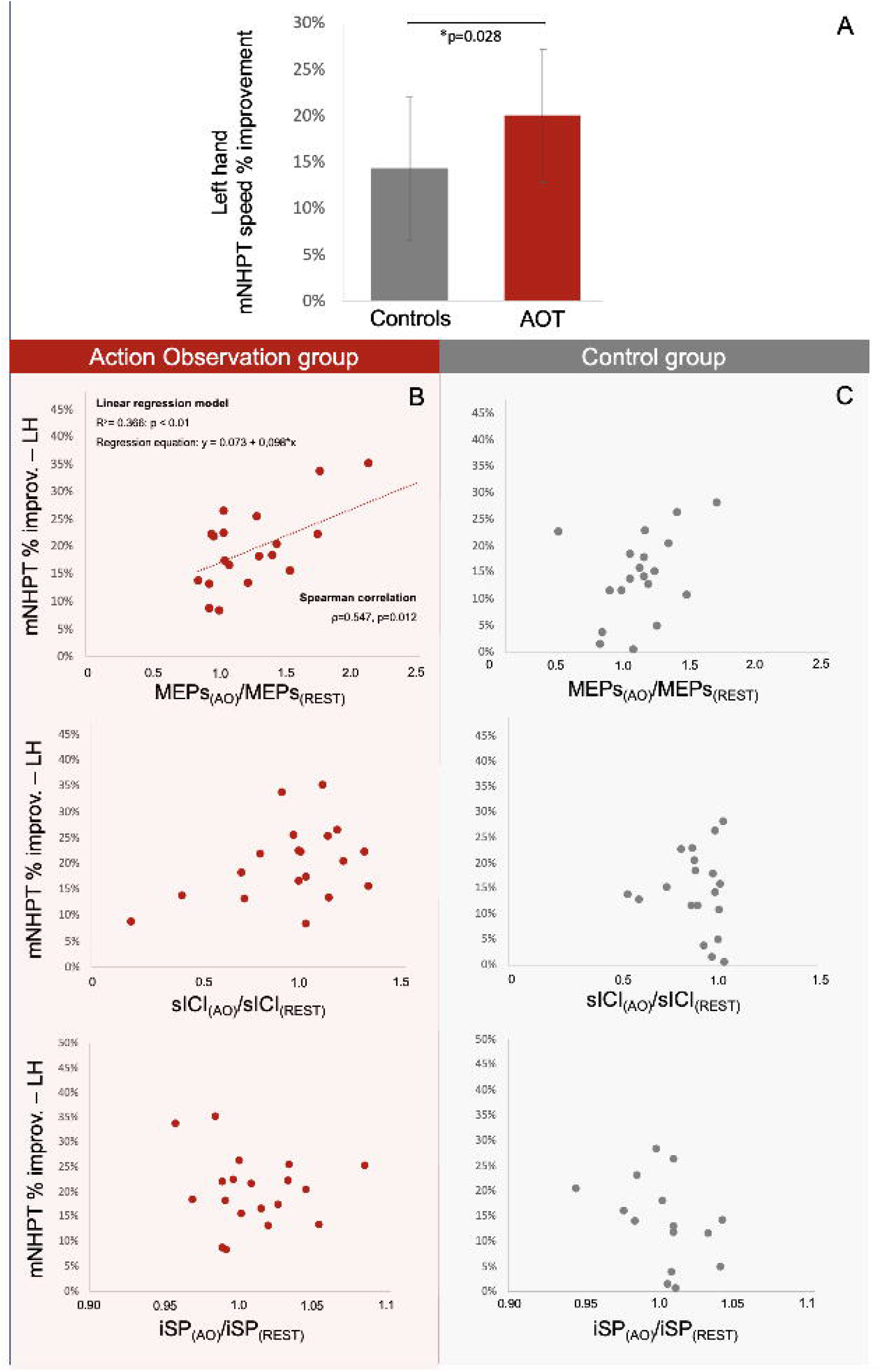

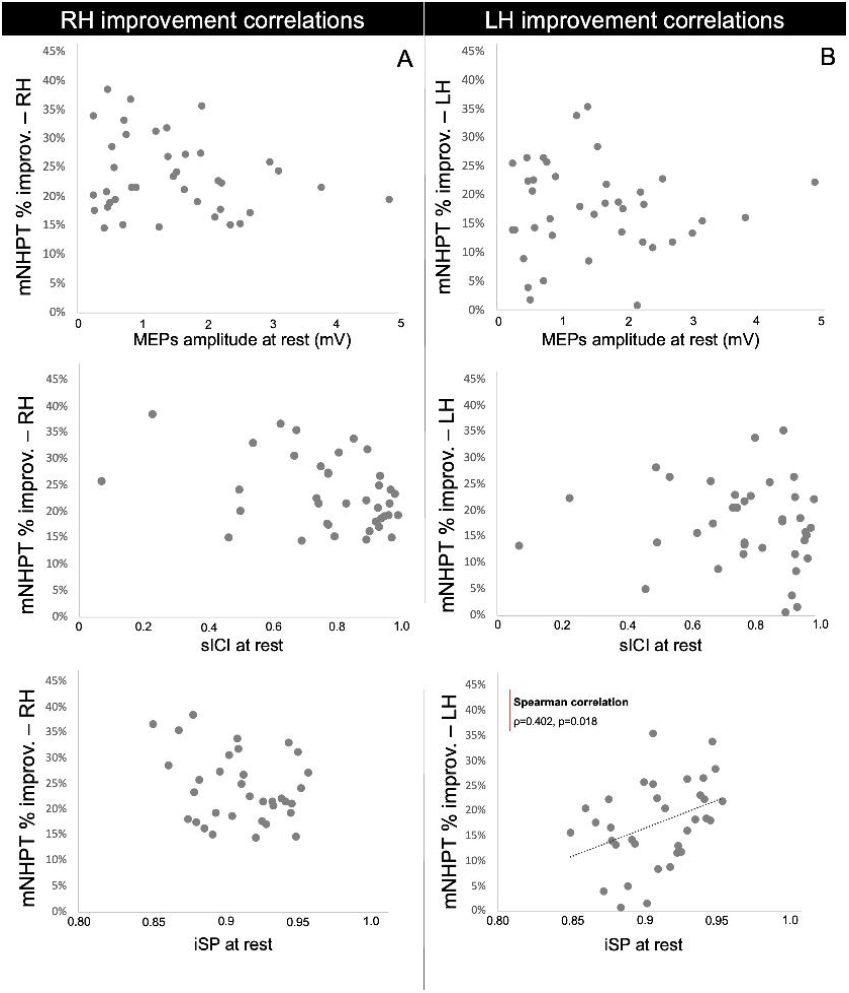

### Neurophysiological predictors of motor improvement

Neither MEPs nor sICI was correlated with the motor improvement of both right- and left-hand performance (see Figure S1). Concerning iSP, no correlation was found with the right-hand performance increase, while a significant correlation was found with the left-hand improvement (ρ = 0.402, p = 0.018).

To investigate whether the TMS features evoked by action observation were associated with the motor improvement promoted by AOT, the MEP amplitude gain in response to action observation (MEPs_(AO)_/MEPs_(REST)_) and the relative increase in intracortical inhibition during action observation (sICI_(AO)_/sICI_(REST)_) were correlated with the improvement in right- and left-hand mNHPT speed, separately for the two groups.

In the AOT group, right-hand improvement was positively correlated with MEPs_(AO)_/MEPs_(REST)_ (ρ = 0.629, p = 0.003) and sICI_(AO)_/sICI_(REST)_ (ρ = 0.733, p<0.001). Linear regression showed that both MEPs_(AO)_/MEPs_(REST)_ (R^2^ = 0.304, p<0.001) and sICI_(AO)_/sICI_(REST)_ (R^2^ = 0.604, p<0.001) constituted significant predictors of the right-hand motor improvement following AOT (see Figure 3, Panel B). The multiple linear regression model confirmed the stronger predictive value of sICI_(AO)_/sICI_(REST)_ in comparison to MEPs_(AO)_/MEPs_(REST)_ (R^2^ = 0.624 vs. R^2^ = 0.339, p<0.001) but also indicated their combination as the best predictor of right-hand motor improvement (R^2^ = 0.680, p<0.001). Bayesian factors confirmed a lower level of evidence for MEPs_(AO)_/MEPs_(REST)_ (BF_1|0_ = 5.64) relative to sICI_(AO)_/sICI_(REST)_ (BF_1|0_ = 312.01) and their combination (BF_1|0_ = 234.33), both indicating a decisive level of evidence [25].

Moving to the left hand, a positive correlation between motor improvement and only MEPs_(AO)_/MEPs_(REST)_ was found (ρ = 0.547, p = 0.012). A subsequent linear regression model identified MEPs_(AO)_/MEPs_(REST)_ as a predictor of motor improvement.(R^2^ = 0.366, p<0.01, see Figure 4, Panel B). Here, the correspondent Bayesian model returned a BF_1|0_ of 5.193, indicating a substantial level of evidence [25] in favor of the alternative hypothesis. It is worth noting that the correlational analyses involving the controls did not return any significant results (see Panel C of Figures 3 and 4), supporting that the AOT motor outcome is specifically associated with the effect of action observation on MEPs and sICI.

### Regression fitting model

Individual data of right-hand performance were fitted with a logarithmic model (*y* = *A* * log(*bx*), where *x* indicates the trial number [see Methods]). Subjects’ curves showed excellent fitting values (all p<0.05), with adjusted R^2^ ranging from 0.605 to 0.960 (mean R^2^ = 0.833). The comparison of *A* coefficients between groups showed higher values in AOT subjects than in controls (t[39] = −3.785; p<0.001; η^2^ = 0.279), supporting that AOT biases the whole motor learning trajectory beyond the single timepoints.

In line with the previous analysis, a linear regression was tested between the estimates of the *A* coefficient and neurophysiological features. Both MEPs_(AO)_/MEPs_(REST)_ (R^2^ = 0.329, p<0.001) and sICI_(AO)_/sICI_(REST)_ (R^2^ = 0.575, p<0.001) were significant predictors of *A*, thus extending the predictive power of such neurophysiological signatures on time-independent AOT outcome.

## Discussion

In this study, we aimed at identifying the neurophysiological signatures explaining motor improvement induced by AOT. We first collected TMS measures assessing the effects of action observation on corticomotor excitability. Then, in a subsequent randomized-controlled experiment, we demonstrated the superiority of AOT relative to motor practice in driving motor learning. Finally, we proved that the modulation of (1) corticospinal excitability and (2) intracortical inhibition induced by action observation successfully predicts individual susceptibility to AOT.

### Action observation effect on corticomotor excitability

One of the simplest ways to probe corticospinal excitability is delivering a single pulse of suprathreshold TMS on a cortical motor map while recording, at the corresponding muscular level, the evoked motor potentials (MEPs), whose amplitude reflects the amount of activated cortical and spinal motor neurons. Consistent with previous reports [6, 7, 12], we found that action observation enhances corticospinal excitability, as measured by MEP amplitude. Three, not mutually exclusive, anatomical models can be adopted to explain such motor-output facilitation. First, the enhancement of primary motor cortex excitability may be driven by excitatory cortico-cortical projections from the premotor [26] and parietal [27] areas. Second, direct [28] descending projections from premotor areas endowed with a mirror mechanism to the spinal cord may increase the pool of recruited spinal motoneurons, resulting in higher MEP amplitude. Third, cortico-striatal neurons endowed with a mirror mechanism may modulate the corticospinal gain, in line with previous findings in animal models [29, 30]; future investigations combining TMS with the administration of pharmacological dopaminergic modulators during action observation will help to validate this hypothesis.

While MEP amplitude is related to the number of corticospinal neurons activated at a given stimulus intensity [31], paired-pulse TMS measure of intracortical inhibition (sICI) reflects the excitability of distinct, low-threshold, GABAergic interneural circuits within the motor cortex [16, 31–34]. Even with a remarkable interindividual variability, we found that action observation, overall, provokes a transient downregulation of corticomotor inhibitory circuits. This result is in line with previous research, where sICI decrease has been related to action observation [7, 35], joint action [9], and action mistake observation [10].

Conversely, we did not find modulation of interhemispheric inhibition during action observation, in partial contrast with a previous report [11]. Methodological differences, such as the use of video clips instead of a live actor, as well as the absence of EMG-based dynamical TMS triggering, could have determined such divergences in results.

It is worth noting that a relevant proportion (35%) of participants showed suppression of MEP amplitude during action observation. Although surprising, this finding is consistent with previous studies [36, 37] and is not incompatible with the neurophysiological models above. Indeed, it could be envisioned a *“behavioral strategy”* view [12, 38, 39] where the excitability of motor pathways is first enhanced by action observation but subsequently suppressed to a greater extent by inhibitory projections from the prefrontal cortex and inferior frontal gyrus [40–42], when subjects requested to volitionally refrain from movement would “repress the urge to act” [38, 39].

Another interpretation is that the decrease in MEP amplitude could reflect an action observation–induced inhibitory activity of interneurons hosted in the primary motor cortex [12]. However, the notion that MEP suppressors do not correspond to sICI enhancers and the absence of a correlation between MEP suppression and sICI during action observation make this latter perspective less likely.

### Electrophysiological predictors of action observation training efficacy

A huge body of research indicates that action observation can help empower existing motor competencies, especially for motor skills requiring fine control [3]. Here, we experimentally confirmed that AOT improves hand motor skills, even in the limb contralateral to that observed and actively practiced. Such an improvement is predicted by the modulations induced by action observation on corticospinal excitability—that is, the greater the MEP facilitation during action observation, the greater the motor skill improvement induced by AOT.

From a neurophysiological perspective, we could speculate that the neural mechanism transiently ignited by action observation could result in neuroplasticity changes, leading to better AOT outcomes [1, 43–45]. Supporting this view, it has been recently demonstrated that the synaptic efficiency potentiation of premotor-to-M1 connections—a key neuroanatomical pathway underlying motor facilitation *via* mirror mechanism [5, 27, 46, 47]—determines the improvement of NHPT performance [23]. An alternative, complementary view is that the repetitive activation of pyramidal tract neurons from premotor areas projecting to the spinal cord may induce spinal plastic changes [48] associated with hand motor control improvement [49]. Interestingly, the predictive role of right-hand muscle facilitation evoked by action observation extends to the left-hand AOT-induced improvement, consistent with the acknowledged bi-hemispheric recruitment of sensorimotor areas during the observation of unimanual actions [50].

Motor tasks’ speed and accuracy depend on the proper selection of which muscles *to move* and which ones *not to move* in each action instant—that is, the appropriate balance between muscle excitation and inhibition [39]. This capacity is even more crucial when dealing with visuomotor tasks [51], precise hand movements [52], and tool use [53]. Moreover, GABAergic cortical activity drives *surround inhibition*, a mechanism that increases the level of segregation of motor activity [54]. TMS measures of intracortical inhibition may constitute a valuable, indirect index of such motor selectivity [51, 54, 55]. We found that the modulation of sICI by action observation largely predicts AOT efficacy, explaining more than 60% of its variance; specifically, subjects with higher increases in intracortical inhibition during action observation showed outperforming AOT learning curves. The evidence in favor of this relationship is strong, more than three hundred times more likely than the *no link* hypothesis.

Recent findings have shown that visuomotor properties in the action observation network might be represented by cell classes that include inhibitory interneurons [56]. We propose that the ability to upregulate such inhibitory circuits in response to action observation may favor the instantiation of inhibitory motor engrams [1], ultimately improving the executive control of the correspondent motor program.

The identification of electrophysiological signatures explaining AOT efficacy may represent the *experimental prelude* to the development of predictive assessments for the selection and identification of the best candidates for AOT in rehabilitative settings and motor training contexts. Extending such knowledge to clinical frameworks would help clinicians to improve the accuracy of prognoses and tune treatment plans, ultimately optimizing patients’ rehabilitation pathways [57]. In this framework, it is worth noting that TMS parameters may be abnormal in several common neurological diseases [17, 57], and future investigations applying our procedures to specific neurological conditions must be envisioned.

## Conclusion

Here, we identified, for the first time, the electrophysiological signatures predicting AOT outcome. Among them, intracortical inhibition modulation plays a major role. We advance that, rather than a volitional “hand-brake” on undesired motor output, the upregulation of such inhibitory mechanisms via action observation may play a key role in the fine-tuning of motor programs, ultimately improving the correspondent performance. Besides its theoretical significance, our study could pave the way for the development of neurophysiological models predicting AOT outcome at the individual level, answering the current need to optimize the rehabilitative pathway of multiple clinical conditions.

## Acknowledgements

We would like to thank Prof. Luca Bonini for his helpful suggestions. The authors confirm that they had no interests which might be perceived as posing a conflict or bias.

## Notes

### Competing Interest Statement

The authors have declared no competing interest.

