## Supplementary material for "The value of corticospinal excitability and intracortical inhibition in predicting motor skill improvement driven by action observation": Figure captions

**Figure 1**

**Study design**

In the first phase (i.e., baseline evaluation), corticospinal excitability modulation by action observation and bilateral hand motor performance were assessed (T0). In the second phase, participants were randomized into two groups. Action observation training subjects were asked to observe a video clip showing a correct execution of the modified version of the Nine Hole Peg Test (mNHPT) and then execute it as quickly and accurately as possible. This observation-execution combination was repeated six consecutive times (T1–T6). The last trial (T6) also included left-hand mNHPT execution. Participants in the control (landscape-observation) group followed the same procedure, except the video clip preceding mNHPT execution depicted an animated lake landscape. mNHPT performance was recorded across T0–T6 timepoints.

Note: TMS, transcranial magnetic stimulation; R, right; L, left.

**Figure 2**

**Effect of action observation on neurophysiological variables.**

Effect of action observation on peak-to-peak motor evoked potential amplitude (Panel A), short-interval intracortical inhibition (Panel B), and transcallosal inhibition (Panel C). Bar charts represent the mean value of neurophysiological variables in the overall population at rest and during action observation.

Note: FDI, first digital interosseous; MEP, motor evoked potential; sICI, short-interval intracortical inhibition; iSP_AMOUNT_, transcallosal inhibition; AO, action observation

**Figure 3**

**Right-hand motor improvement induced by action observation training and neurophysiological predictors of efficacy**

**Panel A.** Right-hand modified version of the Nine Hole Peg Test variations across evaluation timepoints in action observation training (red lines) and control group (gray lines). Single-subject learning trajectories and mean values are represented in thin and thick lines, respectively.

**Panel B.** Scatterplot showing the interplay between right-hand modified version of the Nine Hole Peg Test T0–T6 improvement in the action observation training group and (1) motor evoked potential amplitude gain induced by action observation (top) and (2) intracortical inhibition relative increase during action observation (bottom). Note the significant positive correlations and regressions.

**Panel C.** Scatterplot representing the same variables of Panel B in the control group. Here, no significant correlations were found.

**Figure 4**

**Left-hand motor improvement induced by action observation training and neurophysiological predictors of efficacy**

**Panel A.** Left-hand modified version of the Nine Hole Peg Test T0–T6 changes between two groups.

**Panel B.** Scatterplot showing the interplay between left-hand modified version of the Nine Hole Peg Test T0–T6 improvement in the action observation training group and (1) motor evoked potential amplitude gain induced by action observation, (2) intracortical inhibition relative increase during action observation, and (3) interhemispheric inhibition relative increase during action observation. Motor evoked potential amplitude gain induced by action observation and resultant left-hand motor improvement significantly correlated.

**Panel C.** Scatterplot representing the same variables of Panel B in the control group.

**Supplementary figure 1**

**Neurophysiological basal parameters and motor improvement.**

Scatterplot showing the interplay between right-hand (panel A) and left-hand (panel B) mNHPT T0-T6 improvement in the overall population and: (1) MEPs amplitude at rest (1^st^ row), (2) Intra-cortical inhibition (sICI) at rest (2^nd^ row), (3) Inter-hemispheric inhibition (iSP) at rest (3^rd^ row).
